# Improved Spatial Transcriptomics Clustering with Nested Graph Neural Networks

**DOI:** 10.1101/2025.02.03.636316

**Authors:** Atishay Jain, David H. Laidlaw, Ying Ma, Ritambhara Singh

**Affiliations:** Department of Computer Science, Brown University, 02906, Rhode Island, United States of America; Center for Computational Molecular Biology, Brown University, 02906, Rhode Island, United States of America; Department of Biostatistics, Brown University, 02906, Rhode Island, United States of America

**Keywords:** Graph Neural Networks, Spatial Transcriptomics, Clustering, Machine Learning, Gene Networks, Explainability

## Abstract

We introduce a novel approach, STING, for spatial transcriptomic clustering analysis. Unlike existing state-of-the-art techniques that use graph-based neural networks (GNNs) trained on graphs generated from the spatial proximity of tissue locations (or spots), STING incorporates spot-specific related genes. This feature allows STING to better distinguish between clusters and identify meaningful gene-gene relations for knowledge discovery. It is a nested GNN framework that simultaneously models gene-gene and spatial relations. Using the gene expression, we generate a spot-specific gene-gene co-expression graph. We implement an inner GNN for these graphs to generate embeddings for each location. Next, we utilize these embeddings as features in a sample-wide graph generated using spatial information. We implement an outer GNN for this graph to reconstruct the original gene expression data. Finally, STING is trained end-to-end to generate embeddings that capture gene-gene and spatial information, which we input to a clustering algorithm to produce the spatial clusters. Experiments for 26 samples across 7 datasets and 5 spatial sequencing technologies show that STING outperforms the existing state-of-the-art techniques with a 1.58% to 4.07% improvement in the clustering evaluation metric, thus confirming that integrating gene-gene relation information with the clustering task leads to more informative embeddings and better clusters. Furthermore, experiments on a human breast cancer dataset show that STING identifies relevant genes and gene-gene relations, enabling biological hypothesis generation.

## Introduction

Spatial transcriptomics (ST) has emerged as a potent technology that enables us to measure gene expression in tissues while preserving spatial information. ST has allowed researchers to observe sequencing data along with cell interactions in their spatial environments. Understanding spatial structure is critical in heterogeneous tissues such as brain and cancer since cell function is affected by its environment [26, 38]. Consequently, ST has been used to identify spatially variable genes [7, 1, 39, 21, 43], understand cell-cell communication [36, 4, 20, 15], and cluster cells [40, 41, 24, 12, 42]. In particular, cell clustering is an important step for cell annotation [14, 18], identifying marker genes [16, 22], and understanding disease pathology [8, 5].

Multiple techniques based on hidden Markov random fields (HMRFs) [42, 13] and graph neural networks (GNNs) [24, 41, 44, 12, 16] have been developed to cluster ST data. HMRF methods utilize the gene expression and spatial proximity of spots to cluster them directly. Meanwhile, GNN methods model the ST data as a graph where each node represents a spot, and the edges represent spatial proximity to generate embeddings for the spots. They then input these embeddings into existing unsupervised clustering algorithms such as louvain [2], leiden [31] or mclust [27]. GNN methods have generally shown better clustering performance in multiple benchmarking studies [40, 17]. In particular, GraphST [24] and MuCoST [41] have shown state-of-the-art performance in unsupervised clustering tasks. In addition to GNNs, they utilize contrastive learning to learn better spot representations through spatial modeling.

While all the existing methods perform good clustering through graph-based spatial information modeling, they do not include spot-level gene-gene relations. Since genes interact with other genes differently in distinct cell types [28] and ST clusters have different cell type compositions, we hypothesize that integrating these spot-level gene-gene relations will help improve clustering performance. Furthermore, if these gene-gene relations are explicitly modeled, they can be used to identify important genes and their relations to other genes in a cell-type-specific manner. This will assist with simultaneous biological discovery and improved clustering performance.

We introduce STING (**S**patial **T**ranscriptomics cluster **I**nference with **N**ested **G**NNs), a nested GNN-based framework to integrate spatial information with spot-level gene-gene relations while explicitly modeling the latter. This formulation identifies important genes and gene-gene relations per cell and, by extension, per cell type. To achieve this, we model the spatial information and the gene-gene relations as two different graphs. The spatial information graph is created by treating each spot in the ST data as one node, while the spatial proximity of the nodes defines the edges. Therefore, there is only 1 spatial graph for the entire ST slice/sample. Meanwhile, we generate a gene-gene relation graph for each spot. For these graphs, each node represents a gene, and the edges are determined by their co-expression. Since each spot has its own gene-gene relation graph, one can visualize the entire dataset represented by a graph of graphs, where the outer graph is the spatial graph and all the inner graphs are the spot-level gene-gene relation graphs. As seen in Figure 1, this leads to a nested graph structure. Therefore, STING integrates spot-level gene-gene relations with spatial information to improve clustering performance and explicitly models the gene-gene relations to identify relevant genes and gene-gene relations in specific cell types.

**Fig. 1.**
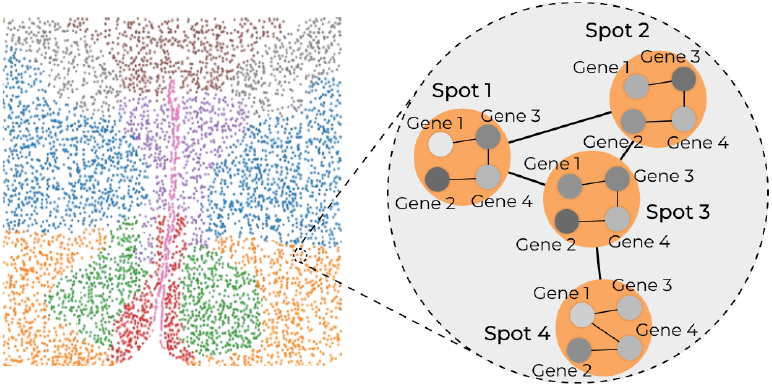
Since each spot contains a graph of genes while also being a node in another graph, we are left with a nested graph structure.

We comprehensively test our framework on 26 samples in 7 datasets across 5 ST technologies with varying spots, clusters, and genes. We show that our framework performs better than state-of-the-art methods GraphST and MuCoST. When compared sample-by-sample, STING has a 1.58% and 4.07% improvement in the clustering performance metric compared to GraphST and MuCoST, respectively. Through an ablation experiment, we also show that the gene-gene relation graphs contribute to the improved performance of STING. On removing all the edges in the gene-gene graphs, there is a 2.74% drop in performance. Finally, experiments on a human breast cancer dataset show that STING can identify important genes and gene-gene relations, thus enabling biological hypothesis generation.

## Materials and Methods

### STING Framework

We introduce a nested GNN framework for unsupervised clustering, as shown in Figure 2. To include the spot-level gene-gene relation information, we first generate gene-gene relation graphs for each spot in the sample.

**Fig. 2.**
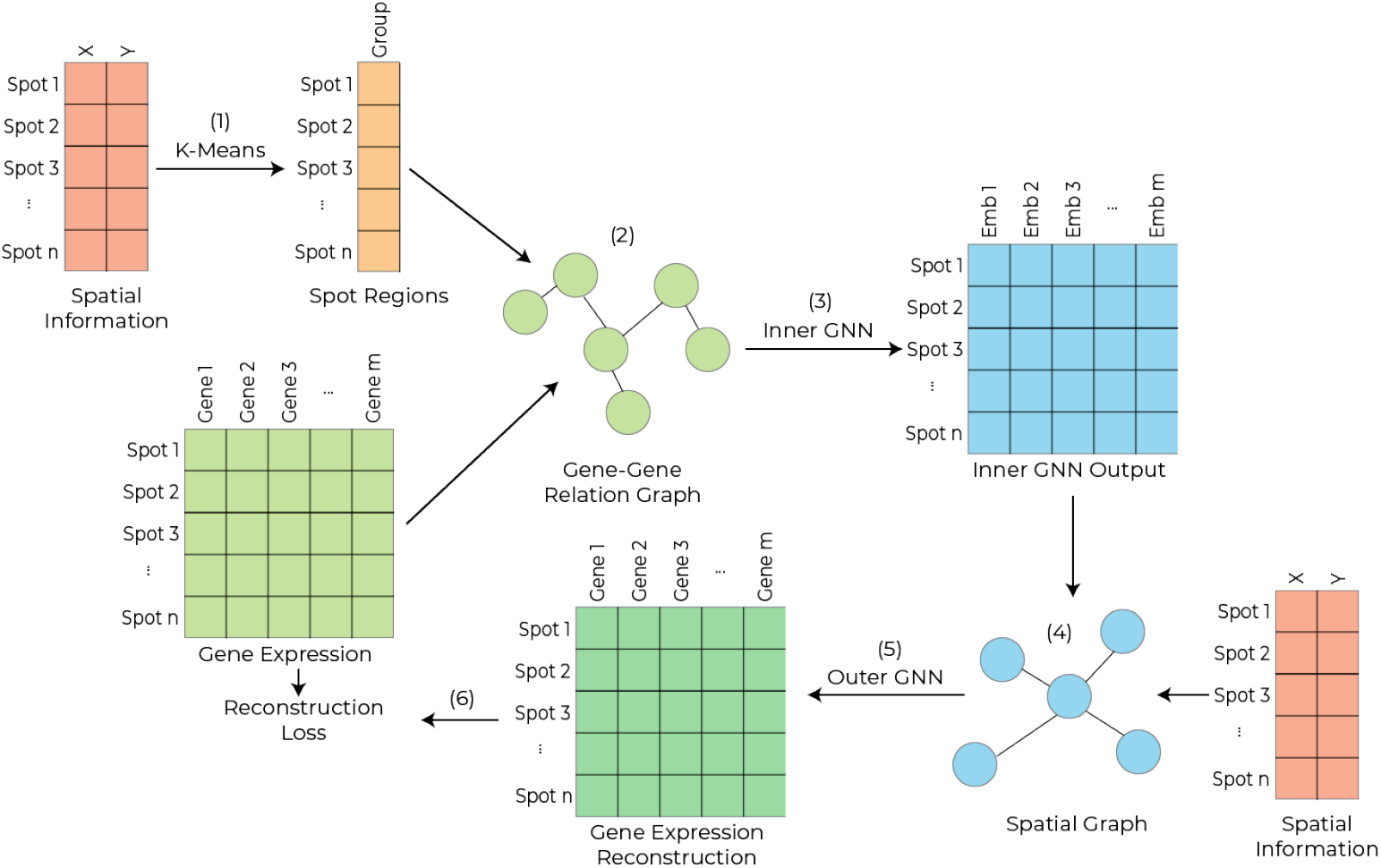
An overview of the STING framework. The overall framework consists of (1) generating spatial regions of spots in the sample. Gene co-expression matrices are generated per region. (2) Spot-specific graphs are generated with edges from the co-expression matrices and features from gene expression inputs and (3) passed through the inner GNN. (4) These inner GNN outputs are used as features in sample-level graphs whose edges are generated using the spatial information of the spots. (5) These graphs are used to train the outer GNN, which aims to (6) reconstruct the original gene expression values.

#### Gene-Gene Relation Graph Generation

Let the 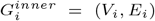 represent the gene-gene relation graph for spot *i*, where *V*_*i*_ is the set of nodes (here, genes), and *E*_*i*_ is the set of edges (here, gene relations). While all the spots share the same genes, *V*_*i*_ differs from spot to spot since we use the gene expression values for the node features. To generate the edge sets *E*_*i*_, we assume that spots of the same cluster are spatially close and that cells of the same cluster have similar gene-gene relations. Based on these assumptions, we generate contiguous regions of spots, which we call spot regions, by applying the k-means algorithm on the spatial coordinates of the spots. Next, for each spot region, we calculate the gene-gene co-expression matrix. Finally, we threshold and binarize the matrix to generate the adjacency matrix for the spot region. Each spot region generally contains homogeneous spots from a cluster, thus retaining cluster-specific knowledge.

If we were to calculate only one gene-gene co-expression matrix from the entire sample, the graphs resulting from such a matrix would be noisy as they would combine information from all clusters and be biased due to potentially disproportionate cell type representations. Our spot region-based approach enables us to create gene-gene relation graphs that are less affected by information in other clusters.

Let 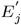 be the edge-set generated from the adjacency matrix for spot region *j*. Then, for all spots *i* in spot region 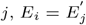.From testing, we noted that STING performed the best with spot regions of size 100 and a threshold calculated from our heuristic that balances the effect of the number of genes and sequencing depth as described in Supplementary Section B.

#### Inner GNN Implementation

We input all the gene-gene relation graphs into our inner GNN. It consists of graph attention network (GAT) v2 layers [3]. We use GATv2 layers since their use of attention scores allows us to identify which gene-gene relations are considered more important by the method. The output of a GATv2 layer is calculated as:

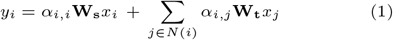

The attention score *α*_*i,j*_ is calculated as:

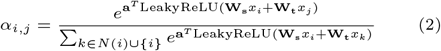

where,

*y*_*i*_ = the output feature vector for node *i*

*x*_*i*_ = the input feature vector for node *i*

**a, W**_**s**_ and **W**_**t**_ = learnable weight parameters

*N* (*i*) = the neighborhood of node *i*

#### Outer GNN Implementation

After passing the gene-gene relation graph through the inner GNN, we get an output graph representation for each spot. This graph representation can be flattened to form a vector which acts as an intermediate embedding for its respective spot. The length of this vector is equal to the number of genes, but the values do not correspond to individual genes. Instead, these values contain joint information about the input gene expression values and the gene-gene relations. Stacking all these vectors results in a *n × g* size matrix where *n* is the number of spots, and *g* is the number of genes.

Despite the inner GNN output matrix not representing the input gene expression matrix, it can be used as an input to existing ST unsupervised clustering algorithms. Therefore, we model our outer GNN on the GraphST method due to its state-of-the-art clustering performance [17, 40]. GraphST is an unsupervised clustering method that utilizes GNNs and contrastive learning. Therefore, we treat the ST data as a graph where each spot is a node, and we use our inner GNN output matrix as the input node feature for each spot. The outer GNN has two primary components. The first component is a GNN-based autoencoder model that aims to reconstruct the input gene expression matrix via a mean-squared error loss:

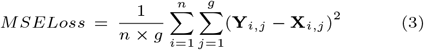

where,

**X** ∈ ℝ^*n×g*^ = input gene expression matrix

**Y** ∈ ℝ^*n×g*^ = output reconstruction of the GNN

*n* = number of spots

*g* = number of genes

The second component is a contrastive learning technique based on Deep Graph Infomax [32]. Given the original graph *G*, the method creates a corrupted/shuffled version of the graph, denoted as *G*^*′*^. With *G* and *G*^*′*^ as inputs, the encoder of the GNN-based autoencoder calculates embedding matrices **Z** and **Z**^*′*^, respectively. To generate pairs for contrastive learning, the outer GNN calculates the readout (*g*_*i*_) of each spot *I* in *G*, representing the spot’s local context. The readout is defined as the sigmoid of the mean of the embeddings of the spot’s neighbors. The embedding of spot *i* (**Z**_*i*_) and its readout *g*_*i*_ together form a positive pair. Similarly, a negative pair is created by calculating the readout of the spots in *G*^*′*^. Specifically, for spot *i*, readout vector 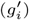 and embedding **Z**^*′*^_*i*_ form the negative pair. Contrastive learning aims to maximize the mutual information of positive pairs and minimize the mutual information of negative pairs. It achieves this through the contrastive loss defined as:

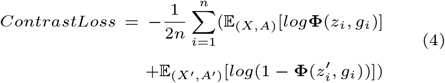

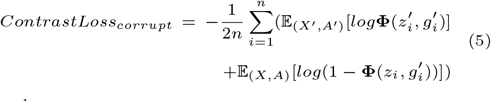

where,

*X, X*^*′*^ ∈ ℝ^*n×g*^ =node features of *G* and *G*^*′*^

*A, A*^*′*^ ∈ *{*0, 1*}*^*n×n*^ =adjacency matrices of *G* and *G*^*′*^

**Φ** =discriminator function that differentiates positive pairs from negative pairs

By combining all the loss functions, we get the final loss for the outer GNN:

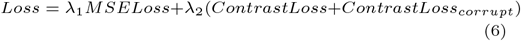

where *λ*_1_ and *λ*_2_ are hyper-parameters used to weight the different types of losses.

Note that we connect the inner and outer GNNs to create one cohesive framework. Therefore, we can train the entire STING framework end-to-end without performing additional calculations in backpropagation. Finally, we use the mclust algorithm [27] on the embeddings generated by the outer GNN to obtain the final clustering.

### Experimental Setup

#### Datasets

We benchmark our clustering performance using 7 datasets from 5 different spatial transcriptomics platforms. All the datasets have manually annotated clusters that we can use as ground truth to test our method. Dataset 1 is a 10x Visium dataset of 12 human dorsolateral prefrontal cortex slices [25]. Dataset 2 is a MERFISH dataset of 5 mouse preoptic hypothalamus samples [26, 23]. Dataset 3 is a BaristaSeq dataset of 3 mouse primary cortex samples [9]. Dataset 4 is a STARmap dataset containing 3 medial prefrontal cortex slices with 166 genes [33, 23]. Dataset 5 is the same STARmap dataset extended to 1020 genes. Dataset 6 is a single osmFISH mouse somatosensory cortex sample [10]. Finally, dataset 7 is a 10x Visium human breast cancer tissue [35]. Table 1 lists the number of samples, clusters, spots, and genes in all the datasets.

**Table 1.** A summary of all datasets used in this work. It lists the number of samples, clusters, spots, and genes in all the datasets.

| Dataset Number | Technology | Sample Count | Cluster Count | Spot Count | Gene Count |
| --- | --- | --- | --- | --- | --- |
| 1 | 10x Visium | 12 | 5, 7 | 3,460 - 4,789 | 33,538 |
| 2 | MERFISH | 5 | 8 | 5,488 - 5,926 | 155 |
| 3 | BaristaSeq | 3 | 6 | 1,525 - 2,042 | 79 |
| 4 | STARmap | 3 | 4 | 1,049 - 1,088 | 166 |
| 5 | STARmap | 1 | 7 | 1207 | 1020 |
| 6 | osmFISH | 1 | 11 | 4,839 | 33 |
| 7 | 10x Visium | 1 | 20 | 3,798 | 36,601 |

For data pre-processing, we use scanpy [34] to filter the genes to the top 3,000 highly variable genes (HVGs) if the dataset has more than 3,000 genes. Next, we normalize the gene expression counts by cell. Finally, we log-transform and then normalize these values to 0 mean and unit variance.

#### Baselines

We compare STING to GraphST [24] due to its consistent and state-of-the-art performance displayed in multiple comprehensive benchmarking papers [40, 17]. We also use a very recent method, MuCoST, as a baseline due to its good reported performance compared to GraphST [41]. While MuCoST is not tested in a comprehensive benchmarking paper, Zhang *et al*. [41] show that MuCoST performs better than GraphST for some datasets.

#### GraphST

GraphST is an unsupervised clustering method that models the spatial transcriptomics data as a graph where each spot is a node, its gene expression values are the node features, and spatially proximal spots are its graph neighbors. As described in the Outer GNN Implementation subsubsection, it uses a GNN-based autoencoder and contrastive learning.

#### MuCoST

MuCoST is another unsupervised clustering method that utilizes GNNs and contrastive learning. It has a very similar model compared to GraphST, with a significant difference being that the positive pair is generated from a graph with edges between the spots calculated using gene expression similarity in addition to spatial distances between them.

#### Hyperparameter Tuning

For GraphST and MuCoST, we use the default hyperparameters set by the papers. Consequently, we use the same hyperparameters as GraphST for our outer GNN. To ensure fairness, we tune the hyperparameters of our inner GNN on only Dataset 1 since both baseline methods have been tested on it. We then fix these hyperparameters when testing on other datasets, such that all the methods use the same setting (pre-fixed hyperparameters) for unseen datasets. We have included the hyperparameter tuning details for our inner GNN in Supplementary Section C.

## Results

### STING outperforms baseline methods for unsupervised clustering

To test STING’s unsupervised clustering performance, we compare it to our chosen baseline methods, GraphST and MuCoST. We test the three methods for 26 samples in the seven datasets using the Adjusted Rand Index (ARI) metric. ARI is a metric commonly used to test unsupervised clustering performance [24, 40, 17] due to its ability to consider the risk of accidental agreement of clusters.

Figure 3 shows that STING has the highest median performance of 0.6622 over all samples, followed by GraphST with a median performance of 0.6554 and MuCoST with a median performance of 0.6484. Figure 4 displays a dataset-wise comparison of the methods, validating that STING performs better in most datasets and ST platforms. The differing relative performance of STING may be attributed to the sequencing depths of these datasets. This correlation is observed in scatter plots in Supplementary Section A, which show the percentage difference in ARI performance between STING and the baselines with respect to the average sequencing depth of a dataset. Finally, we compare the performances sample-by-sample and note that STING has a mean ARI improvement of 1.58% and 4.07% over GraphST and MuCoST, respectively. A comparison of the methods for all individual samples is available in Supplementary Section D.

**Fig. 3.**
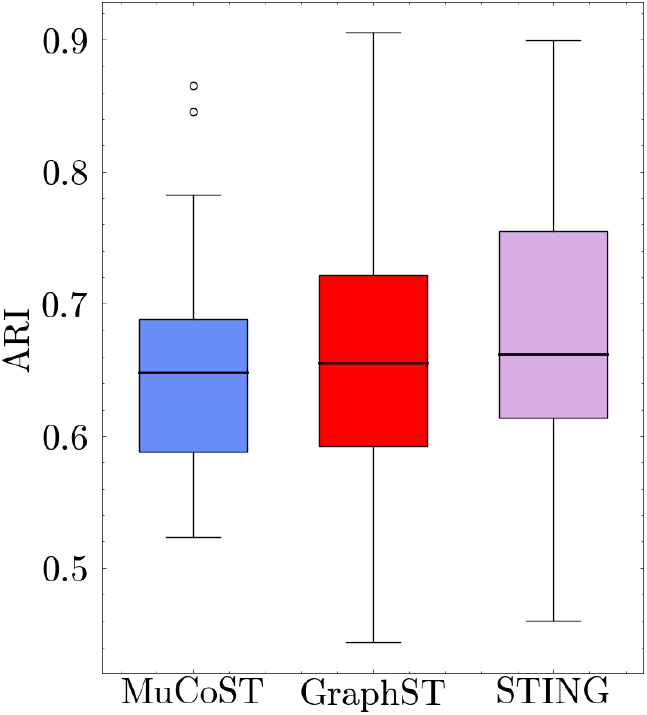
A quantitative comparison of all methods over all samples shows that STING demonstrates the best clustering performance.

**Fig. 4.**
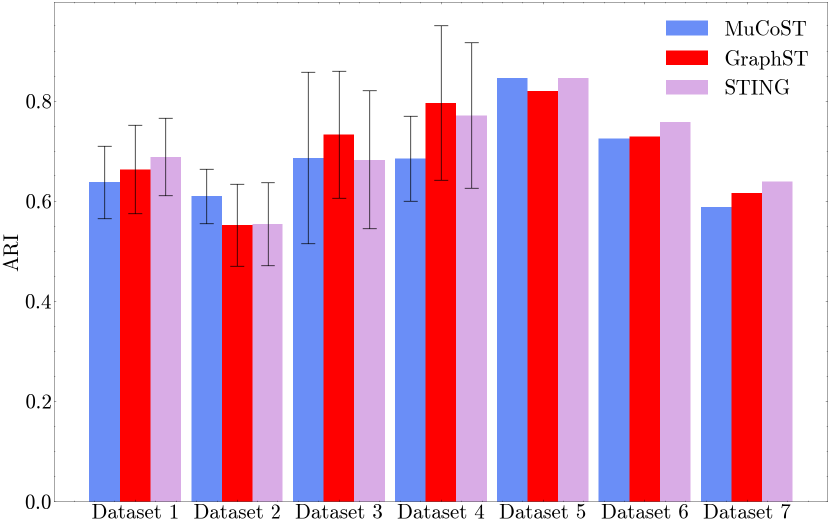
A quantitative comparison of all methods over the seven datasets show that STING has better overall performance on most datasets and ST platforms. The bar plots represent the mean performance on the corresponding dataset, while the error-bars represent the standard deviation.

Qualitatively, Figure 5 shows that STING’s clustering outputs are spatially coherent. In contrast, GraphST and MuCoST sometimes display artifacts where one cluster is spread over the sample, which is undesirable. The areas in solid rectangles of GraphST’s and MuCoST’s outputs contain examples of such artifacts. For example, in panel a of Figure 5, both MuCoST and GraphST split cluster 8 into multiple clusters all over the sample. Similarly, in panel b, MuCoST splits clusters 3, 4, 9, and 10, and GraphST splits clusters 1, 4, 7, and 9. While STING does split clusters 1 and 9, it does so to a much lesser extent. Due to this splitting, GraphST and MuCoST fail to properly identify the ground truth cluster ‘Layer 5’, while STING is able to do so. In general, the visual examples show agreement with the quantitative results.

**Fig. 5.**
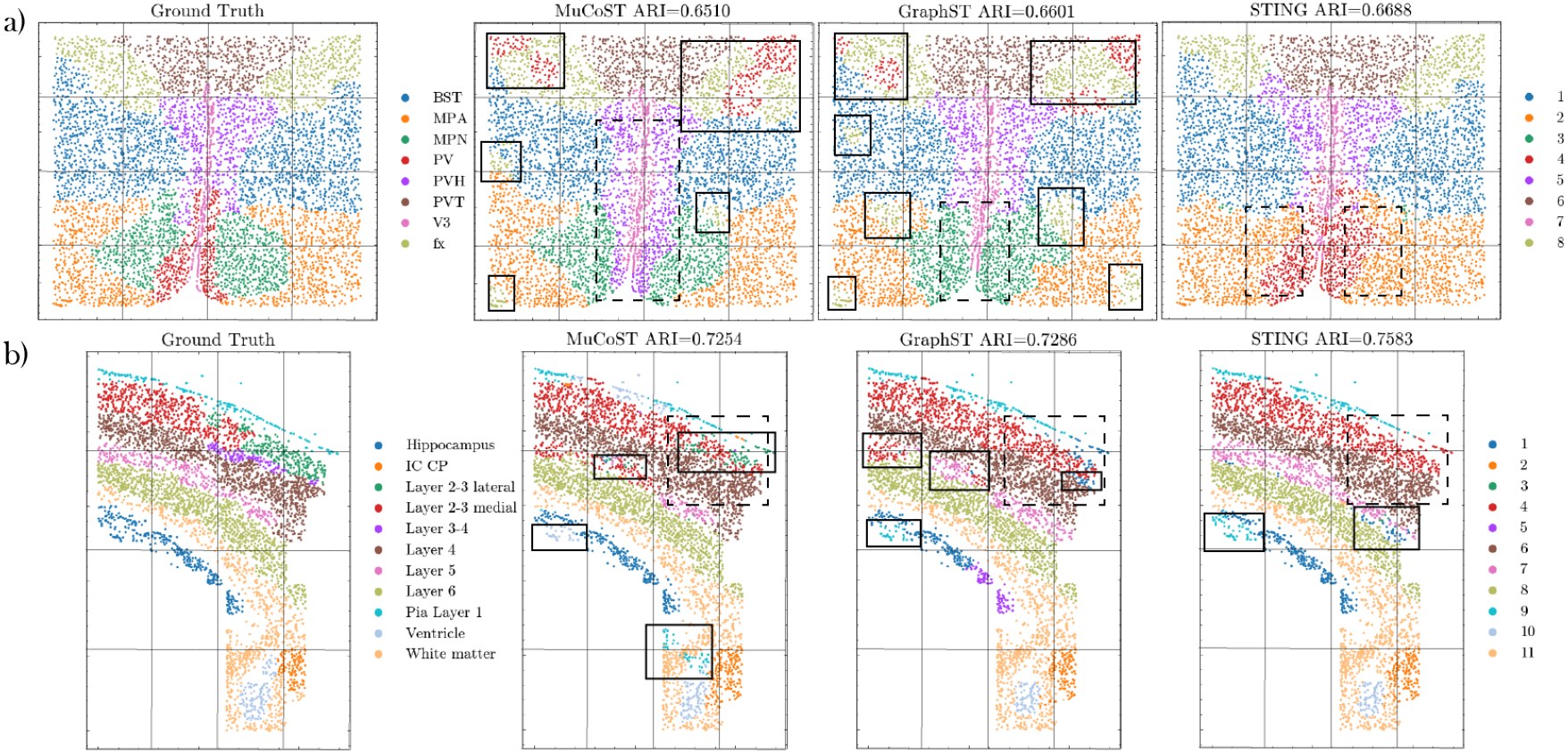
A qualitative comparison shows that STING mitigates the undesirable spatial incoherence of clusters. Clusters within the solid rectangles are examples of clusters broken into multiple parts even where not desired. Meanwhile, regions in dashed rectangles are examples of ground truth clusters not identified by the methods.

We also note that all the methods tend to miss certain ground truth clusters. The dashed rectangles in Figure 5 represent such areas. For example, in panel a, GraphST and STING cannot separate the ground truth cluster ‘MPN’ from its neighboring clusters ‘PV’ and ‘MPA’, while MuCoST merges the ‘PV’ and ‘PVH’ clusters. Similarly, in panel b, none of the methods identify ‘Layer 2-3 lateral’ or ‘Layer 3-4’. This observation could be attributed to the fact that these clusters are generally harder to separate since they are a mix of other layers.

### Ablation studies show that gene-gene relation graphs improve clustering performance

To test whether the gene-gene relation graphs contribute to STING’s performance, we design an experiment where we remove all the edges of the spot-level gene-gene relation graphs before inputting them to STING. Since all other components of the ablated method are the same, the only difference in performance is due to the gene-gene relation graphs.

We compare all 26 samples in the 7 datasets and observe that the ablated STING method underperforms compared to STING (Figure 6). The ablated STING method shows a median ARI performance of 0.6280, while STING has a median performance of 0.6622. Furthermore, a sample-by-sample comparison shows a 2.74% drop in ARI when we remove the gene-gene relation edges. A comparison of the methods over all individual samples is available in Supplementary Section D.

**Fig. 6.**
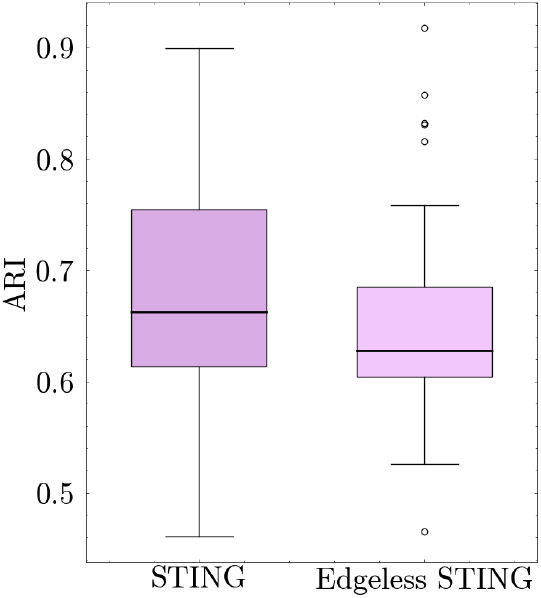
Removing the edges from the gene-gene relation graphs leads to a quantitative decline in clustering performance, thus showing the graphs’ importance.

### STING’s attention scores identify important genes and gene-gene relations

Finally, we test STING’s ability to determine important genes and gene-gene relations via its attention scores. Since there is a lack of methods to validate all the important genes and gene-gene relations, we conduct two experiments utilizing the human breast cancer dataset (dataset 7). This is due to the large volume of human breast cancer research and relevant databases. Furthermore, identifying important genes and gene-gene relations in breast cancer can lead to potential therapeutic gene perturbation targets.

First, we select key genes from the GNN methods and identify pathways among the selected genes. Next, we rank important gene-gene relations and compare them to the similarity of somatic mutations of genes in the COSMIC database [29] given the assumption that there is some correlation between gene-gene co-expression and co-occurrence of gene mutations [19, 11]. We use the attention scores from the first layer of STING’s inner GNN for both experiments. Furthermore, we combine all the clusters in STING’s output representing ductal carcinoma in situ (DCIS) and invasive ductal carcinoma (IDC) clusters in the ground truth and treat them as one cluster *C*. We combine the clusters to minimize heterogeneity within breast cancer and allow us to compare our results to existing standard databases.

We obtain the attention scores (**A**) in the form of a three-dimensional array of size (*n × g × g*), where n is the number of spots ∈ *C* (=1,970), and g is the number of genes (=3,000).

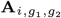 represents the attention score from gene *g*_1_ to gene *g*_2_ in spot *i*. It is also important to note that not every gene-gene relation edge is included in every spot’s graph due to the thresholding step during graph creation. We do not include those gene pairs while calculating average scores for any gene or gene-gene pair.

In the first experiment, we test whether the set of important genes determined by STING are biologically more relevant to the cluster than those obtained from GraphST. To determine the top genes in STING, we calculate the average attention score for every gene and rank the genes based on this average. Then, we choose the top genes that cumulatively contribute to 10% of the sum of these averages to identify pathways for the cluster. We use a percentile-based number to compare different methods, and 10% selects enough genes to identify pathways reliably. For the top genes in GraphST, we use GNNExplainer [37] since GraphST does not include an innate method to identify top genes like STING. GNNExplainer is a widely used method to explain GNN predictions. It identifies sub-graphs, nodes, and features important to the GNN’s prediction. From

GNNExplainer, we can obtain an importance score for each gene in every spot. Like STING, here we calculate the average score for all the genes over all the spots in *C* and choose the top genes that cumulatively account for 10% of the average scores. Since the primary differences between STING and GraphST are the spot-level gene-gene relation graphs and their corresponding attention scores, a better gene selection by STING implies that the attention scores learn cluster-specific biologically relevant knowledge. Thus, the attention scores can assist in discovering genes important to clusters.

We obtain 66 genes for STING and 86 genes for GraphST from the experiment. For each set of genes, we use the STRING database [30] to identify important pathways. For STING, the database recognizes 58 genes and identifies the KEGG pathway ‘Pathways in Cancer’ with the highest number of genes (8). For GraphST, the database identifies 82 genes. It labels 6 genes according to the KEGG pathway ‘MicroRNAs in cancer’. However, the database also labels the ‘Huntington disease’ pathway for 7 genes, the ‘Alzheimer disease’ pathway for 7 genes, the ‘Parkinson disease’ pathway for 6 genes, and the ‘Prion disease’ for 6 genes. GraphST may have selected some of these brain disease-related genes since they are part of the reactome pathways ‘Cell Cycle’ and ‘Developmental Biology’. Therefore, the database picks up multiple pathways for GraphST, where only one is relevant to the cell type. Meanwhile, the pathway labeled for the highest number of genes for STING is relevant to the cell type. Therefore, the cells identified by STING are less ambiguous and more relevant to the cluster, suggesting better discovery of genes relevant to specific cell types.

In the second experiment, we test whether the gene-gene relation ranking obtained from STING contains biologically relevant information. We cannot use GraphST as a baseline in this experiment since it does not model gene-gene relations. Since the nodes in GraphST’s input graphs represent spots and edges represent interactions between spots, the explanation method cannot be used to identify important gene-gene relations.

Since the attention scores can represent any relation between genes, we do not have an existing ground truth to validate these scores. Therefore, we use similarities in breast cancer somatic mutations between genes to test whether the attention scores learn any biologically relevant information. If the ranking of the scores shows some biological relevance, we can use STING to identify important gene-gene interactions for different cell types.

For the experiment, we calculate the average attention score for every gene-gene pair across all samples and rank the pairs based on the average. Gene pairs with an average score of 0 are discarded since they were filtered out when generating the gene-gene relation graphs. We then use the COSMIC database to generate a somatic mutation similarity score for every pair that was not discarded. For every gene, we obtain sample IDs in the COSMIC database in which the gene shows a mutation. We define the mutation similarity score as a simple Jaccard score. Between two genes *g*1 and *g*2 the similarity score is defined as:

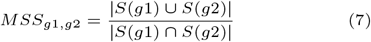

where,

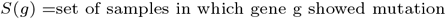

Since none of the baseline methods can score gene-gene relations, we compare STING’s gene-gene pair ranking to a ranking determined by the gene-gene co-expression values. STING uses gene-gene co-expression values to generate the gene-gene relation graphs. Therefore, if its ranking performs better than the gene-gene co-expression-based ranking, we can conclude that STING can extract additional biologically relevant information due to its joint training.

First, we compare both rankings’ normalized discounted cumulative gain (NDCG). NDCG is a commonly used metric to measure and compare ranking quality and is defined as the discounted cumulative gain (DCG) of the given ranking divided by the DCG of the ideal ranking. DCG is defined as:

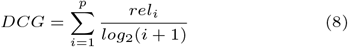

where,

*p* = number of pairs

*i* is based on method ranking

*rel*_*i*_ = ground truth score of pair *i* (here, score from COSMIC)

Over 5 runs, STING’s ranking has a mean NDCG of 0.8013 with a standard deviation of 0.005. It is consistently higher than the gene-gene co-expression-based ranking’s NDCG of 0.793.

To further compare the two rankings, we calculate the percentage of top-k pairs common to the ground truth and the method rankings. For k, we choose 0.1%, 0.2%, 0.5%, 1%, 2%, 5%, and 10% of the total number of pairs in the ranking. As seen in Figure 7, STING outperforms the gene-gene co-expression-based ranking.

**Fig. 7.**
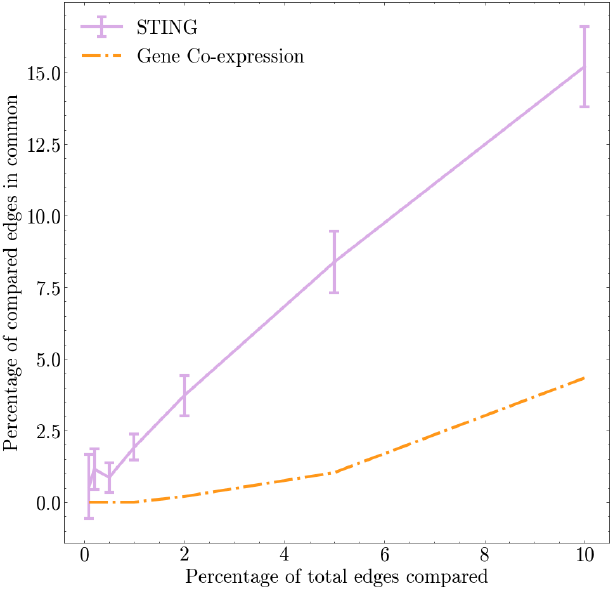
When averaged over 5 runs, the gene pairs from STING show a better somatic ranking when compared to the gene-gene co-expression-based ranking. The line plots represent the mean performance, while the error bars represent the standard deviation.

We conclude that STING can extract some information on somatic mutation similarities between genes from both comparisons. This is despite the method having no prior knowledge of somatic mutations and the gene-gene relation graphs being solely calculated based on gene-gene co-expression in that single sample. Therefore, the experiments show that STING’s attention scores learn additional biologically relevant information that could help identify important genes and gene-gene relations in various cell types.

## Discussion

This paper presents a nested GNN framework, STING, that performs unsupervised clustering on ST data. The main advantage of STING over existing clustering methods is that it explicitly models spot-level gene-gene relations. The inclusion of gene-gene relations adds valuable information to the embeddings, leading to better clustering performance. Experiments on multiple datasets display STING’s performance improvement over state-of-the-art methods. Moreover, using datasets from different ST platforms also highlights STING’s generalizability. Furthermore, an ablation study proves that the gene-gene relation graphs lead to better clustering. These experiments make STING an attractive choice for unsupervised ST clustering.

Since these gene-gene relations are explicitly modeled in STING, we can directly identify important genes and gene-gene relations in clusters. This ability is another advantage of STING over existing methods. Since other methods do not explicitly model gene-gene relations, they cannot identify important gene-gene relations at all. While we can identify important genes using these methods, we must rely on external explanation methods. These explanation methods may not accurately capture what the method is learning, and different methods could give different results. These varying results could lead to confusion and the inaccurate identification of genes.

Additionally, two experiments on a human breast cancer dataset show that the attention scores in STING can learn biologically relevant information. The most important genes obtained from these scores can identify less ambiguous pathways than those obtained by GraphST. Furthermore, the gene-gene pair ranking obtained from the attention scores can learn somatic mutation similarities despite being given no relevant prior knowledge. This suggests that STING may be a viable and helpful tool for simultaneous biological discovery. While modeling gene-gene relations leads to a few benefits, it comes with the limitation of higher computational resource requirements. Since each spot adds a graph whose size increases with the number of genes, the memory requirements of STING scale with the number of spots and genes. One 10x Visium human dorsolateral prefrontal cortex sample consists of 4,789 spots and 3,000 genes. Running STING with this sample requires 23 GB of memory and around 10 minutes to run 900 epochs. In comparison, GraphST requires only 1 GB and 15 seconds to run 600 epochs. With advances in ST technology, samples with a higher number of spots are being generated [6], possibly leading to a memory bottleneck. These memory requirements could be mitigated by reducing the number of genes through HVG selection or reducing the number of spots by dividing the sample into smaller chunks. However, these techniques could lead to lower performance, resulting in a trade-off between performance and resources.

Finally, while describing our framework, we mentioned that the outer GNN of the method was GraphST. We chose GraphST for its best overall clustering performance. However, our framework does not require GraphST as its outer GNN. The outer GNN can be replaced with any method that uses a gene expression matrix as the input. This can lead to the possibility of extending STING’s longevity by using a new and better clustering method as the outer GNN. However, we have not tested this possibility, which remains an open research question.

## Code and Data Availability

The code is publicly available at https://github.com/rsinghlab/STING. The datasets are available on a site hosted by Yuan *et al*. [40] at http://sdmbench.drai.cn/.

