## Supplementary File for "Improved Spatial Transcriptomics Clustering with Nested Graph Neural Networks"

#### A - Preliminary Investigation of Relative Performance of STING and Dataset Sequencing Depth

During experimentation, we noticed that STING's performance relative to the baseline models varied from dataset to dataset. On further investigation into the properties of these datasets, we identified that there was a possible correlation between the sequencing depth of the datasets and STING's relative performance. This result is displayed in two figures that show the relation between the mean percentage difference in ARI score between STING and the baseline methods per dataset and the sequencing depth of the corresponding dataset. Table S1 lists the average sequencing depth of the 7 datasets. Figure S1 shows the relation of STING's relative performance to GraphST, while Figure S2 exhibits the relation of STING's relative performance to MuCoST.

Table S1: The average sequencing depth of all datasets.

| Dataset Number | Average Sequencing Depth |
|---|---|
| 1 | 3,452,034 |
| 2 | 1,400,651 |
| 3 | 83,162.67 |
| 4 | 549,764.7 |
| 5 | 483,149 |
| 6 | 25,782,188 |
| 7 | 18,017,760 |

#### B - Heuristic Selection for Gene-Gene Relation Graphs

To investigate the ideal number of edges for the gene-gene relation graphs, we tested multiple STING models with an average node degree (average number of neighbors of a node) of 1,3,5 and 7. These models showed varying performance over the different datasets. Therefore, we decided to develop a heuristic node degree. In Supplementary Section A, we identified that sequencing depth was a factor in STING performance. Therefore, we plotted the effect of sequencing depth on the ideal node degree, as seen in Figure S3. While most datasets seemed to follow the same trend, Dataset 6 seemed to be a major outlier. Since Dataset 6 had the fewest genes among all the datasets, we investigated whether the number of genes affected the ideal node degree. As exhibited in Figure S4, there seemed to be another trend where an increasing gene count led to an increasing ideal node degree. This trend explained the outlier noted in Figure S3 as the genes limit the effect of sequencing depth on the average degree. After performing non-linear regression with the sequencing depth and gene count as variables, and the ideal node degree as the target, we arrived at a heuristic where:

$$d = 0.35 \times (\frac{s}{100000})^{0.5} \times (ln(g-30) - 1) \tag{1}$$

where, $d$ is the ideal average node degree, $s$ is the sequencing depth, and $g$ is the number of genes.

Since memory requirements increase with average node degree, we also capped the average degree at 7.

### C - Hyperparameter Tuning for Inner GNN

For all the inner GNN, we employed a grid hyperparameter search. The values that we searched over are shown in Table S2

Table S2: The range of values used for each hyperparameter during hyperparameter grid search. The bold numbers represent the final value chosen for that hyperparameter.

| Hyperparameter | Values |
|---|---|
| Number of GAT layers | 1, **2**, 4 |
| Hidden size of GAT layers | **1**, 2, 4 |
| Learning rate | $10^{-2}$, $\mathbf{10^{-3}}$, $10^{-4}$ |

### D - Sample-wise Comparison of STING to Baseline and Ablation Methods

Figure S5 is a sample-wise comparison of the ARI performance of GraphST, MuCoST, STING, and the ablated model of STING. Similarly, Figure S6 is also a sample-wise comparison. However, in this figure, the bars representing a baseline are the differences in ARI score between STING and the corresponding baseline method. Table S3 lists which sample belongs to which dataset.

Table S3: Samples in each dataset.

| Dataset Number | Sample Names |
|---|---|
| 1 | 151507, 151508, 151509, 151510, 151669, 151670, 151671, 151672, 151673,151674, 151675, 151676 |
| 2 | MER4, MER9, MER14, MER19, MER24 |
| 3 | BaristaSeq1, BaristaSeq2, BaristaSeq3 |
| 4 | STARBZ5, STARBZ9, STARBZ14 |
| 5 | STARBY3 |
| 6 | osmfish |
| 7 | HBC |

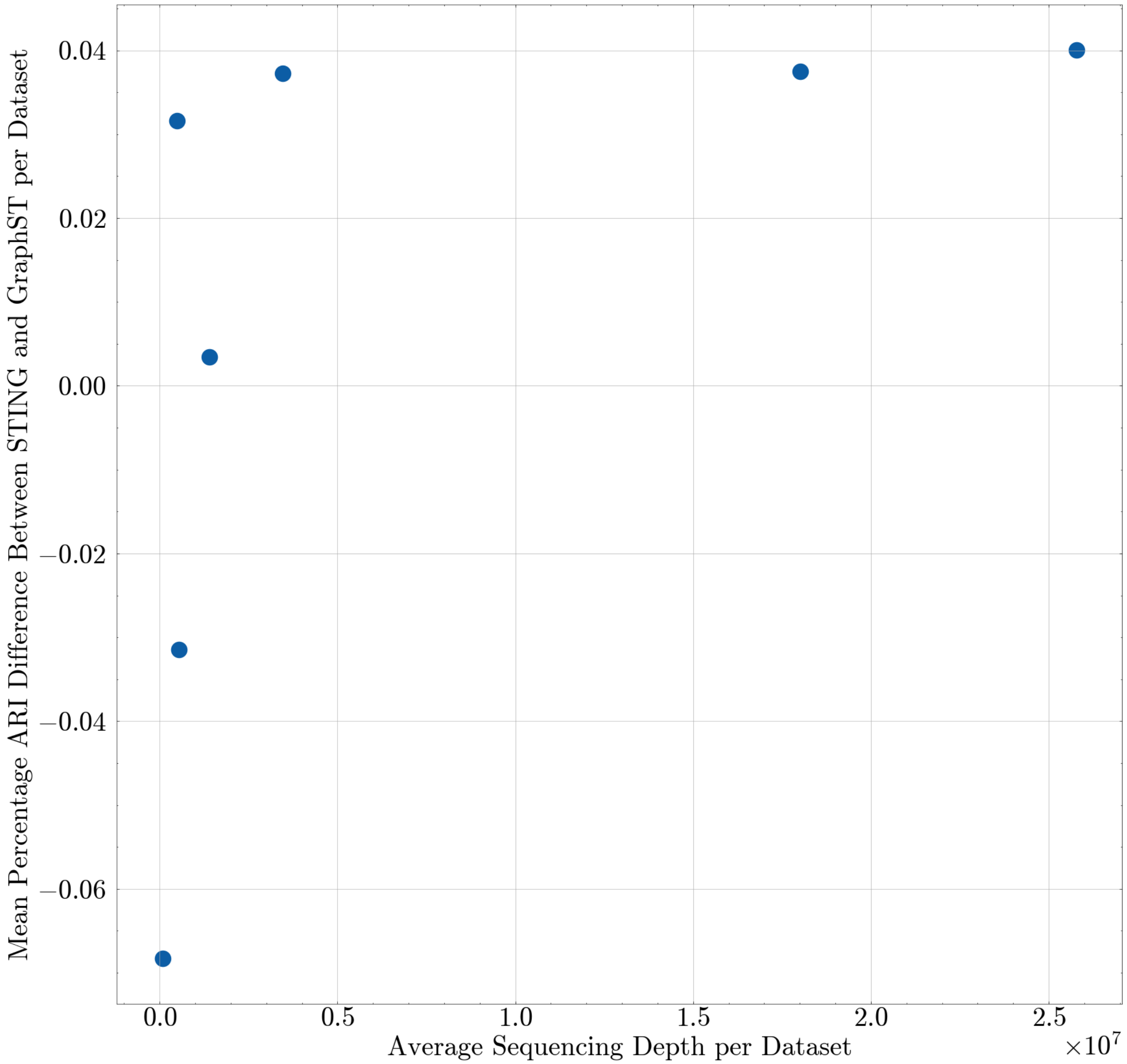


Figure S1: This figure is a scatter plot between the mean percentage difference in ARI score between STING and GraphST in each dataset and the sequencing depth of the corresponding dataset. We can see that there is a general trend where a higher sequencing depth leads to a better relative STING performance.

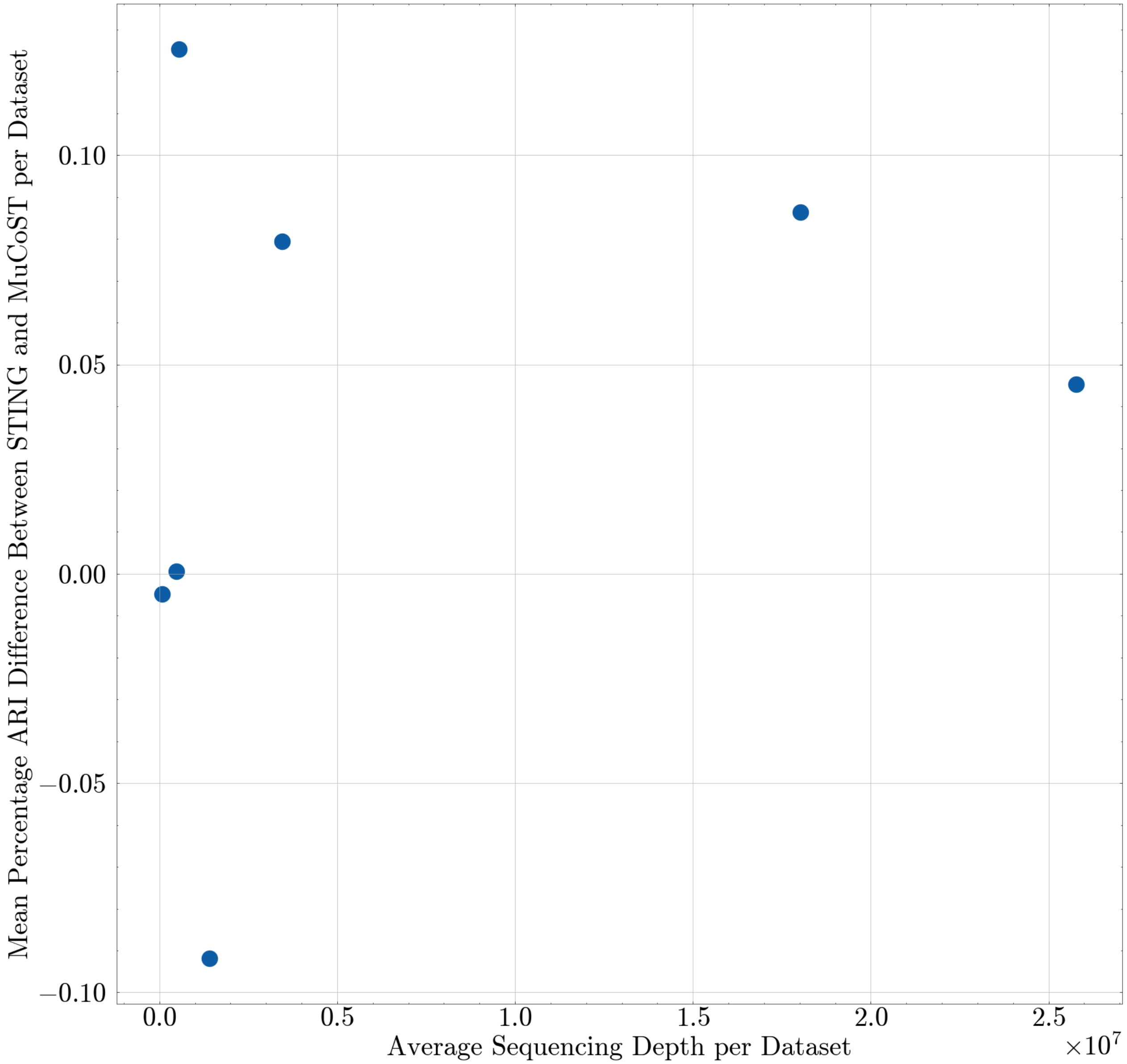


Figure S2: This figure is a scatter plot between the mean percentage difference in ARI score between STING and MuCoST in each dataset and the sequencing depth of the corresponding dataset. We can see that there is a general trend where a higher sequencing depth leads to a better relative STING performance. The trend here is slightly less clear compared to the relative performance of STING and GraphST.

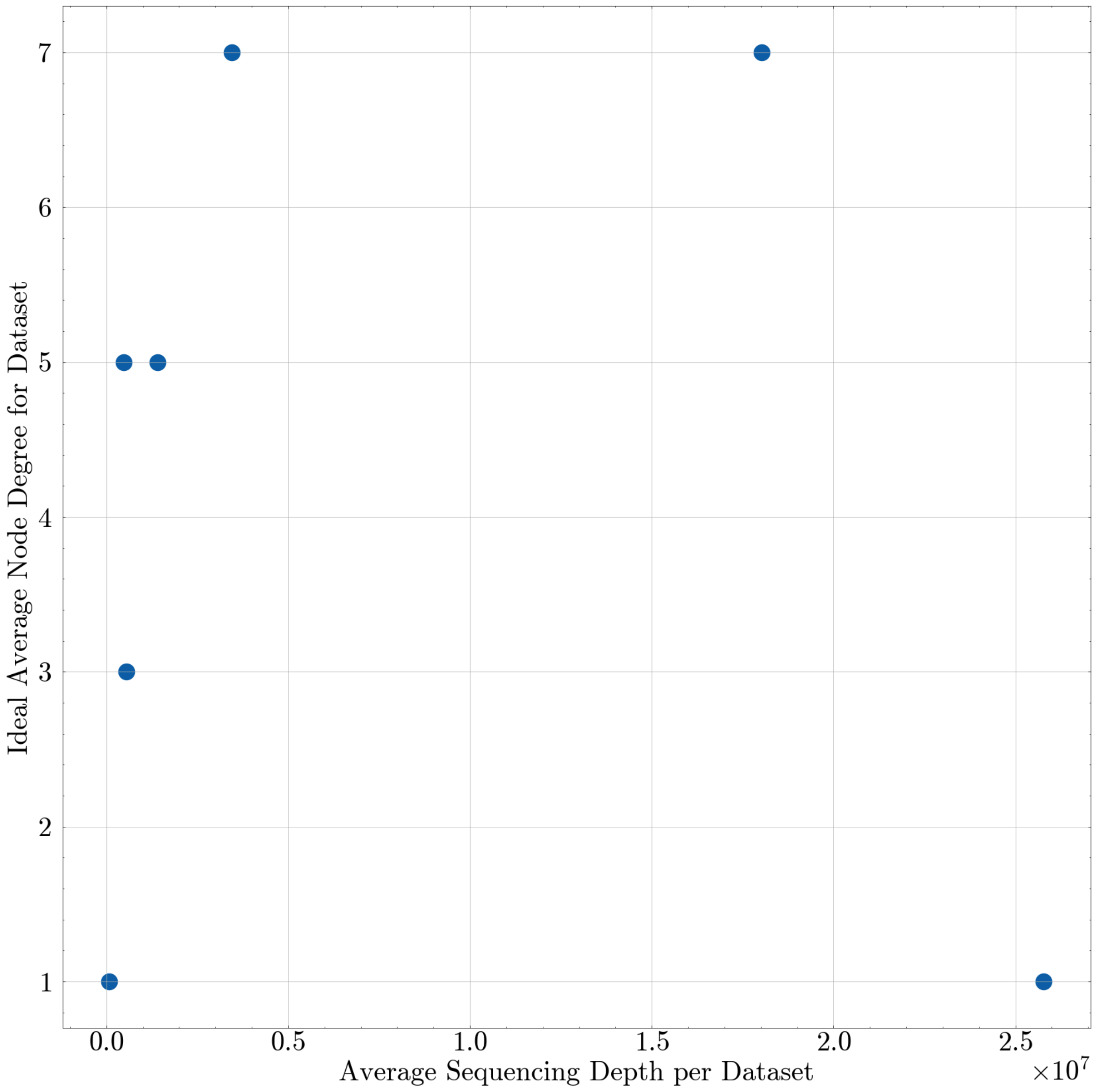


Figure S3: This figure is a plot representing the relation between sequencing depth and ideal average node degree. We can see that an increasing depth leads to a higher degree, with one major outlier (Dataset 6).

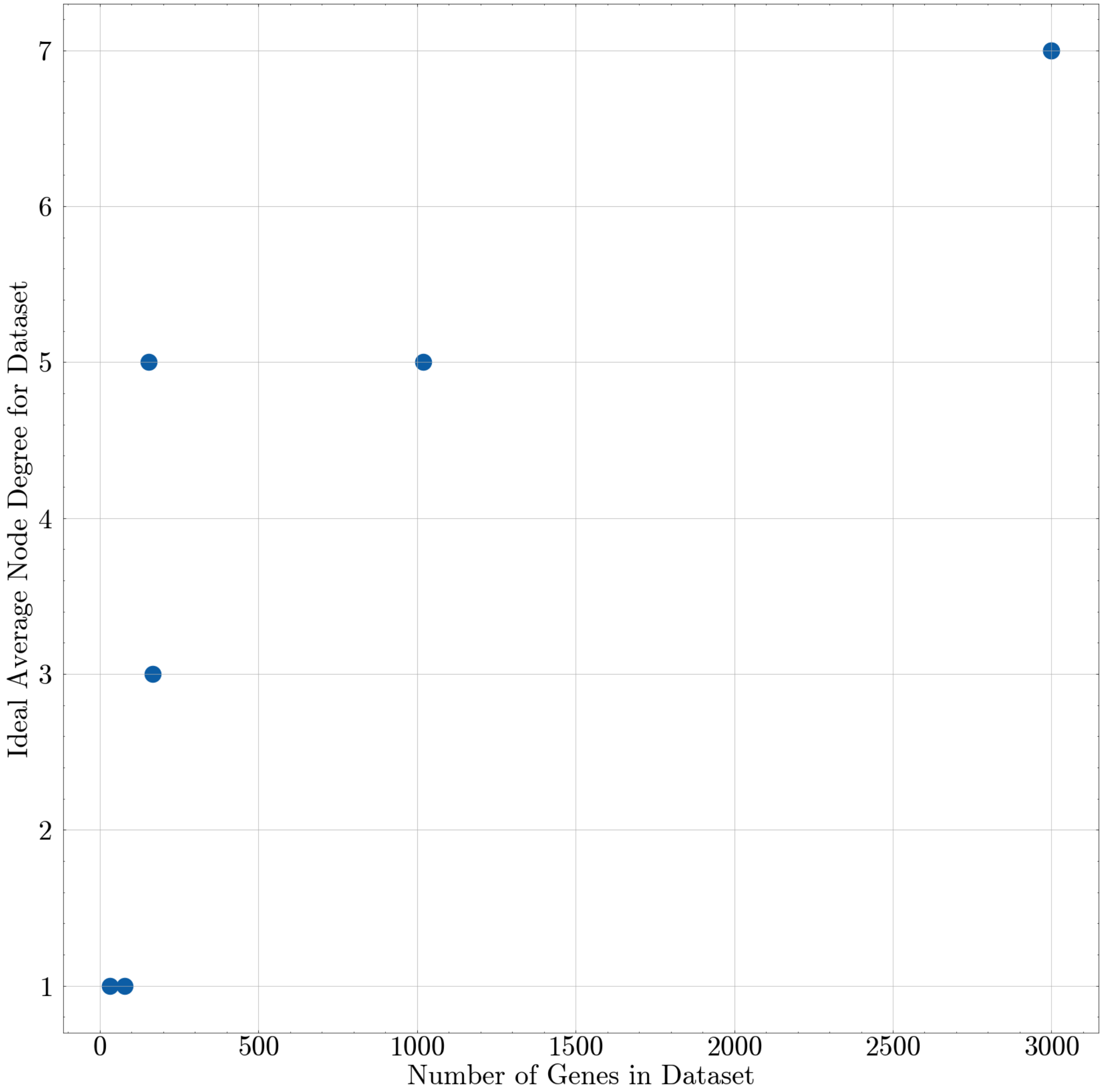


Figure S4: This figure is a plot representing the relation between the number of genes and ideal average node degree. We note that an increasing gene counts generally leads to a higher degree.

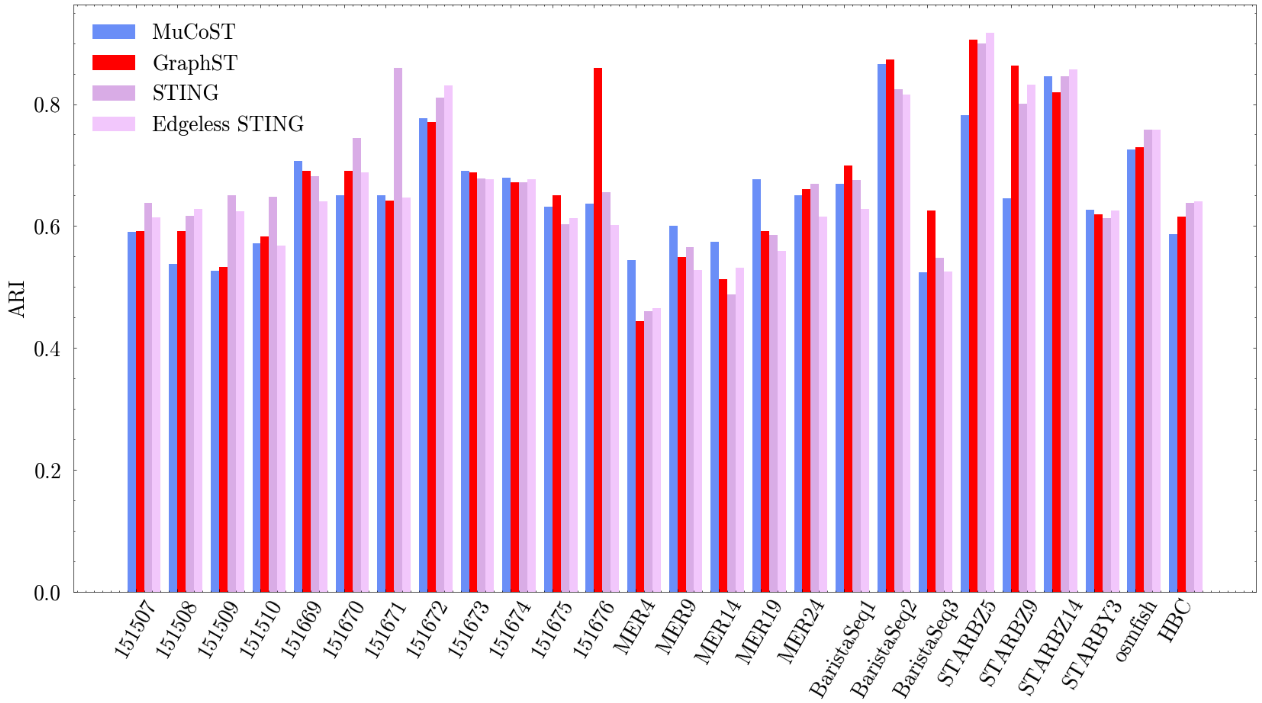


Figure S5: This figure is a sample-wise ARI comparison of GraphST, MuCoST, STING, and the ablated STING model.

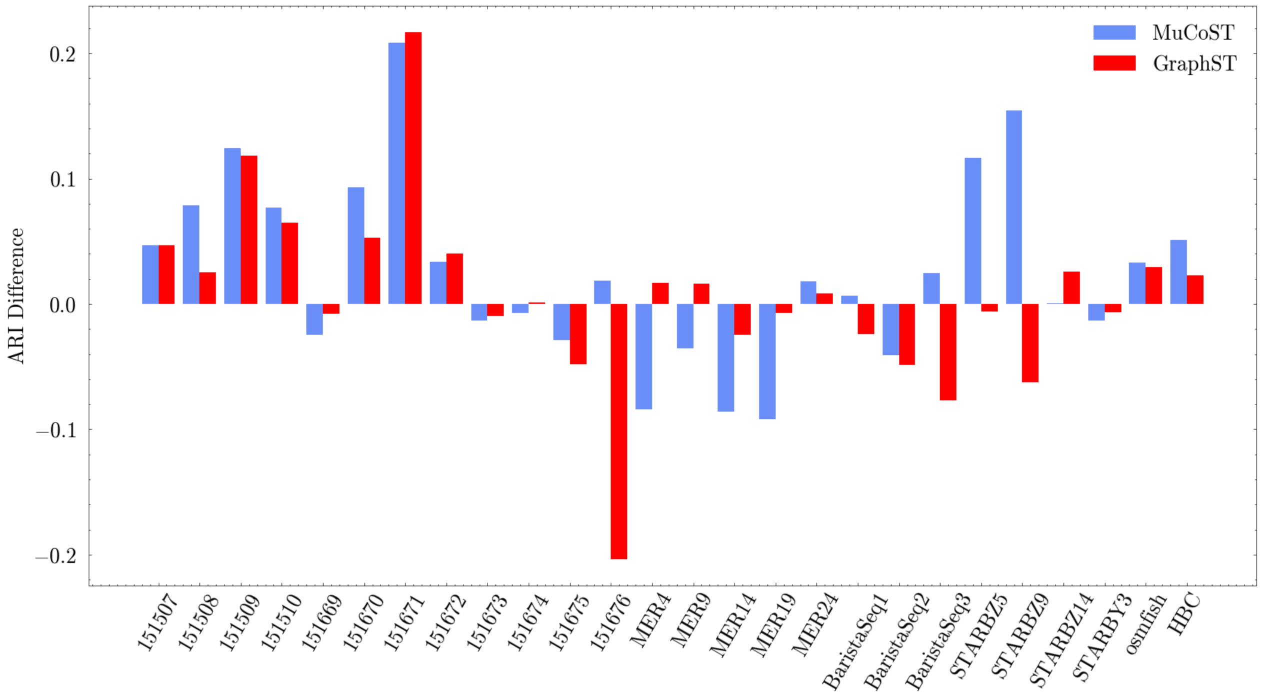


Figure S6: This figure is another sample-wise comparison of GraphST, MuCoST, and STING. Here, the bars represent the ARI score difference between STING and the baseline methods labeled in the figure.
